# The Little Transib That Could – and Could Not: Contrasting Invasion Outcomes within a Novel DDE DNA Transposon Lineage

**DOI:** 10.64898/2026.08.26.747230

**Authors:** Artem Ilin, Mattias Mannervik

## Abstract

Transposable-element (TE) abundance can vary substantially among populations, yet population-specific differences in TE content remain incompletely characterized. Here, we compared known TE families across long-read genome assemblies representing geographically and historically distinct *Drosophila melanogaster* populations. Several genomes showed pronounced strain-specific expansions, including an exceptional increase in copies of the element historically annotated as *Hopper* in A6-Wild5B. Investigation of this expansion revealed a previously uncharacterized full-length autonomous element encoding a 648-amino-acid transposase. We named this element *Nozomi* and its non-autonomous derivative *Kodama*, corresponding to the published *Hopper* consensus. Protein-sequence, phylogenetic and structural analyses placed *Nozomi* within the *Transib* superfamily and showed close correspondence between the predicted *Nozomi* transposome and the experimentally determined *Helicoverpa zea Transib* strand-transfer complex. Autonomous *Nozomi* copies were restricted to a small number of *D. melanogaster* genomes, where they were associated with extensive but separate expansions of *Kodama*. All *Kodama* elements carried the same precise 1,380-bp internal deletion, with breakpoint microhomology suggesting an alternative end-joining-related origin. Comparative searches identified a broader group of related *Transib* elements with contrasting invasion histories. Unlike the restricted *Nozomi* distribution, *Hayabusa* showed minimal sequence divergence, limited structural decay and broad distribution across the *Drosophila* suzukii and montium groups, consistent with a recent, highly successful horizontal invasion. Thus, closely related *Transib* elements can follow markedly different trajectories after horizontal acquisition: *Nozomi* remained restricted while driving local amplification of a shorter non-autonomous derivative, whereas *Hayabusa* spread broadly across species.

## Introduction

Transposable elements (TEs) are widespread mobile genetic elements that occupy substantial fractions of eukaryotic genomes. Although uncontrolled transpositional activity is often associated with deleterious effects on the host, including disruption of genes and regulatory elements^1–3^, chromosomal rearrangements and ectopic recombination^4–8^, TEs have also acted as major drivers of genome evolution over longer evolutionary timescales. TEs are typically maintained within host lineages by vertical inheritance, which requires their activity or retention in germline genomes. However, vertical transmission alone cannot explain the sudden appearance of closely related TE families in distantly related and reproductively isolated species. Such cases are instead interpreted as horizontal transfer (HT) of transposable elements, which is currently the only well-supported mechanism by which an established TE family can enter a new host lineage^9^.

Several essential biological mechanisms are thought to have emerged through TE domestication or exaptation^10,11^. A prominent example is the RAG1/2 recombinase, which is essential for V(D)J recombination and therefore for the adaptive immune system of jawed vertebrates. RAG1 shows clear evolutionary affinity to *Transib*-like DNA transposases, and the discovery of active ProtoRAG elements in lancelets provided direct support for a transposon origin of the RAG recombination machinery^12–14^.

*Transib* elements, named after the Trans-Siberian Express, constitute a comparatively poorly characterized superfamily of eukaryotic cut-and-paste DNA transposons, originally identified from degenerated copies in insect genomes^13,15^. Described *Transib* members encode a DDE/D transposase and are bounded by terminal inverted repeats, with canonical *Transib* insertions typically generating 5-bp target-site duplications. Biochemical and structural characterization of *HzTransib* from *Helicoverpa zea* demonstrated RAG-like DNA cleavage, hairpin formation and strand transfer, establishing *Transib* as an important model for the evolutionary origin of RAG- mediated recombination^14,16,17^. However, the biology of intact *Transib* elements remains poorly understood, particularly with respect to their structural variation, taxonomic distribution and population-level dynamics.

*Drosophila melanogaster* is one of the best metazoan models for TE research. Its genome contains more than one hundred TE families representing both class I retrotransposons and class II DNA transposons^18–20^. In *Drosophila*, TEs have been linked to changes in endogenous gene expression, the evolution of new host functions through domestication, and repeated HT events between species^21^. Population-genomic and historical studies have shown that TE dynamics in *Drosophila* are highly heterogeneous: TE families differ in abundance and transposition rate among species and populations, and multiple families have undergone distinct invasion waves over historical time^22,23^.

Despite more than four decades of TE research in *D. melanogaster*, much of our understanding of population-level TE variation has been derived from cytological assays, family-specific molecular approaches and, more recently, short-read sequencing^23–26^. These studies established extensive variation in TE abundance and insertion frequencies among populations, but repetitive sequence and structural heterogeneity limit the resolution of short-read approaches. Population-scale long-read assemblies now provide direct access to individual TE insertions and their internal structure, revealing a substantial fraction of insertions and structural variants that remain unresolved in short-read datasets^27^.

Here, we used published long-read genome assemblies from geographically and historically distinct *Drosophila melanogaster* populations to identify strain-specific expansions of individual TE families. This analysis led to the discovery of a previously uncharacterized full-length autonomous form of the element historically annotated as *Hopper*^28^. Because *Hopper* is also the name of an unrelated hAT-family DNA transposon from *Bactrocera dorsalis,* we designate this autonomous *Transib*-superfamily element ***Nozomi*** and its centrally deleted 1.4 kb non-autonomous derivative, corresponding to the published *Hopper* consensus, ***Kodama***. We subsequently searched for related elements across *Drosophila* and other dipterans, identifying a broader group of closely related *Transib* elements with notably different patterns of genomic expansion, structural deterioration and horizontal spread.

These comparisons revealed two particularly striking features of this lineage. First, closely related elements showed contrasting invasion outcomes: autonomous *Nozomi* remained restricted to a small number of *D. melanogaster* genomes, whereas the related *Hayabusa* element showed evidence of extensive horizontal spread across species. Second, both the *Nozomi*/*Kodama* and *Transib-6* showed pronounced amplification of internally deleted non-autonomous derivatives of approximately 1,400 bp. Together, these observations identify a previously unrecognized *Transib* lineage characterized by heterogeneous invasion dynamics and a recurring enrichment of shortened non-autonomous forms.

## Results

### Population-scale analysis of TE insertions reveals strain specific expansions of several TE families

Transposable element (TE) abundance can vary substantially among populations, yet the frequency and scale of exceptional, genome-specific expansions remain poorly understood. To examine this variation at insertion-level resolution, we analyzed annotated TE families across two complementary panels of long-read *Drosophila melanogaster* genome assemblies spanning broad geographic and temporal ranges. The first one consisted of 14 founder-strain assemblies from the Drosophila Synthetic Population Resource (DSPR), representing globally sampled strains collected in the 1920s–1960s and subsequently maintained as laboratory stocks^29,30^. The second consisted of 32 assemblies generated by the DrosEU long-read genome project, including 24 wild European populations caught between 2014 and 2017, and 8 Drosophila Reference Genomic Panel lines derived from a single population from Raleigh, USA, sampled in late 1990s-early 2000s^27^.

Comparison of the historical stock-derived DSPR genomes with more recently sampled natural populations provided a first view of TE dynamics across these panels. As expected, we detected strong differences for recently invading TE families, including the *P-element* and *Transib1*^21^ (Fig. S1A,B).

Focusing on the DSPR founder genome panel, we compared putatively recent, genome-specific TE insertions by retaining only copies close to the length of their corresponding consensus, excluding nested insertions and removing loci shared with other analyzed genomes. Several individual genomes contained unusually high numbers of insertions from particular TE families (Fig. 1A). The endogenous retroviral element *roo* was a notable exception, with large numbers of genome-specific insertions across all analyzed genomes (Fig. 1A).

**Figure 1.**
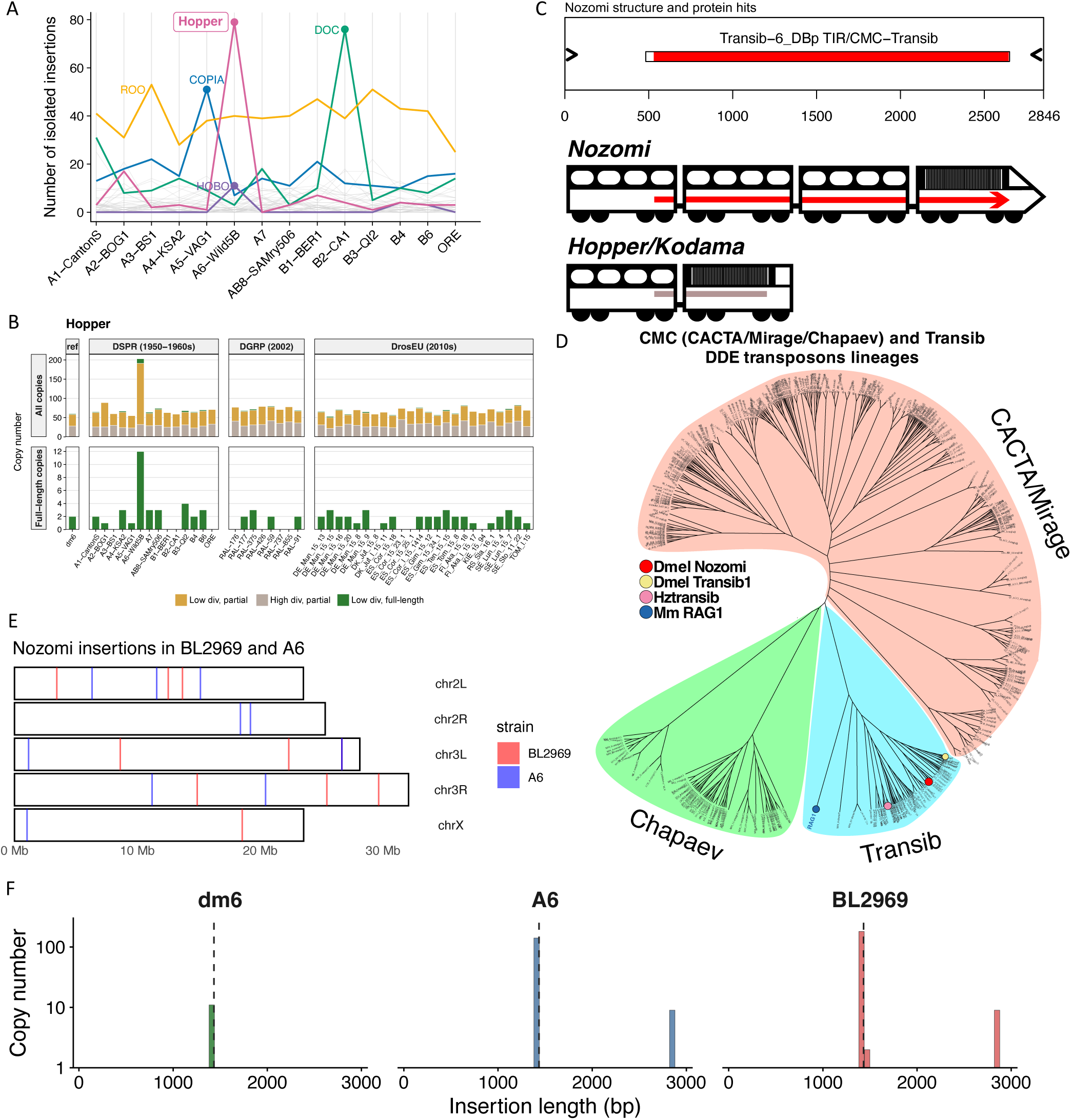
Identification of *Nozomi*, an autonomous DNA transposable element related to the *Kodama* element, previously annotated as *Hopper*. (A) Copy numbers of genome-specific, isolated, long (relative to consensus) insertions of selected transposable element families across the Drosophila Synthetic Population Resource (DSPR) founder genome panel. (B) Copy-number comparison of all *Hopper* insertions regardless of length, uniqueness or proximity to other insertions across DSPR and DrosEU long-read genome panels. Top: insertions classified as diverged partial, non-diverged partial and non-diverged full-length copies. Bottom: non-diverged full-length insertions only. Insertions were classified as diverged when they differed from the consensus sequence by more than 10%. (C) Structure of the *Nozomi* mobile element, showing two 37 bp terminal inverted repeats (TIR; black arrowheads) and an ORF encoding a protein with high similarity to the *Transib-6* DNA transposon from *Drosophila bipectinata*. *Kodama* elements are approximately two times shorter than *Nozomi* and are non-autonomous, as depicted on the scheme. (D) Phylogenetic tree of closely related CMC (CACTA/Mirage/Chapaev) and *Transib* DNA transposons. *Nozomi* and *Transib1* from *D. melanogaster*, *HzTransib* from *H. zea,* and RAG1 protein from *M. musculus* are shown as coloured circles. (E) Karyoplots showing the coordinates of *Nozomi* insertions in the *D. melanogaster* A6 and BL2969 genomes after liftover to the dm6 genome assembly. (F) Size distributions of *Nozomi* and *Kodama* insertions containing both TIRs in the dm6, A6 and BL2969 genomes. Dashed lines show the median size for all insertions in a genome.

The remaining expansions were largely restricted to individual genomes and involved multiple TE classes. The A5-VAG1 genome, derived from flies collected in Athens, Greece, in 1965, showed an expansion of the LTR retrotransposon *copia*. The B2-CA1 genome, derived from a population collected in Cape Town, South Africa, in 1954, showed an increased copy number of the LINE-like retrotransposon *Doc*. In contrast, the A6-Wild5B genome, derived from flies collected at Red Top Mountain, Georgia, USA, in 1966, showed an extensive expansion of the Terminal Inverted Repeat (TIR) DNA transposon *Hopper*. Comparison across the combined DSPR and DrosEU panels showed that this pronounced *Hopper* expansion was unique to A6- Wild5B (Fig. 1B). Most *Hopper* insertions in A6 were located in distal intronic or intergenic regions, but over 20% occurred within 1 kb of gene transcription start sites (TSSs), a pattern not observed for *Hopper* insertions in the dm6 reference genome (Fig. S1C).

Although *Hopper* represented the most pronounced TE expansion in A6-Wild5B, this genome also contained an elevated number of *hobo* insertions (Fig. 1A). The increased abundance of these DNA transposons coincided with occurrence of structural differences, some of which were flanked by *Hopper* or *hobo* insertions. For example, comparison of the A6-Wild5B genome with both the A1-Canton-S and dm6 assemblies revealed an inversion in A6 that is associated with a *Hopper* insertion on one flank and no detectable TE on the other (Fig. S1D).

The expansion of different TE families in the A5, A6 and B2 genomes suggests that these strains depart from the low-copy, quasi-equilibrium pattern expected under simple transposition– selection balance^3,31^. The timing of these expansions remains unresolved, including whether they occurred before or after the strains were established as laboratory stocks. Nevertheless, their restriction to individual genomes reveals substantial heterogeneity in the recent evolutionary histories of TE families within *D. melanogaster*.

### Identification of *Nozomi*, the autonomous partner of *Hopper*

The increased abundance of *Hopper* insertions in the A6 genome was unexpected because *Hopper* had previously been described as a non-autonomous DNA transposon^15,32^. Consistent with this annotation, inspection of the published *Hopper* consensus sequence^15,19^ revealed no open reading frame (ORF) capable of encoding a full-length transposase or any protein. Inspection of individual *Hopper* BLAST hits in the A6 genome revealed approximately two-fold longer insertions that retained strong similarity to the 5′ and 3′ regions of the published *Hopper* consensus but contained an additional internal sequence absent from the short consensus and not detectably homologous to other annotated TEs (Fig. S2A). Extraction of the full sequence revealed an ORF encoding a 648 amino acid protein. Searches against transposable-element protein databases identified *Transib-6* from *Drosophila bipectinata* as the closest match, classifying it as a *Transib*-superfamily DNA transposon (Fig. 1C). This classification is consistent with earlier work that linked the short *Hopper* sequence to *Transib*-superfamily elements, particularly *Transib1*, and proposed that autonomous copies may not have been retained in the *D. melanogaster* reference genome^15^. Because the name *hopper* had previously been also assigned to an unrelated, but more thoroughly described hAT-family transposon from *Bactrocera dorsalis*^33,34^, and to distinguish the autonomous and non-autonomous forms of the *D. melanogaster* element, we designate the newly identified full-length element ***Nozomi*** and the previously described non-autonomous form ***Kodama*** (Fig. 1C). The names *Nozomi* and *Kodama* refer to the same 500-series Shinkansen trainsets, which originally operated as 16-car *Nozomi* formations and were subsequently reconfigured as 8-car *Kodama* formations. This nomenclature reflects the derivation of the shorter non-autonomous *Kodama* element from the full-length autonomous *Nozomi* element through an approximately two-fold reduction in length. Notably, despite the clear similarity to *Transib*-family transposases, the predicted *Nozomi* transposase produced no significant conserved-domain hits in InterProScan with all available member databases used.

To further establish the evolutionary placement of *Nozomi*, we constructed a maximum-likelihood phylogeny using representative DDE transposases from the related *CACTA–Mirage– Chapaev* (*CMC*) and *Transib* clades obtained from the RepeatMasker protein database. The *Nozomi* transposase grouped within the *Transib* clade, in agreement with the protein-similarity searches and with the previously proposed relationship between non-autonomous *Hopper/Kodama* and *Transib* elements (Fig. 1D). Thus, phylogenetic analysis placed the newly identified *Nozomi* transposase within the lineage expected from the sequence characteristics of the previously described short element.

### Distribution of *Nozomi* and *Kodama* insertions in *Drosophila melanogaster genomes*

We identified 12 *Nozomi* copies in the A6 genome. Three of these corresponded to degenerated and divergent copies also present in the dm6 reference genome at the same chr3L locus. In dm6, this locus contained two long *Nozomi* insertions. All degenerated copies lacked the first 8 nt of the 5′ terminal inverted repeat (TIR), which is expected to disrupt their transpositional potential. By contrast, the remaining nine copies in A6 were distinct from the degenerated chr3L copies but were nearly identical to each other, differing only by up to 2 nt in the length of the 3′ poly(dA) region (Fig. S2B).

To determine whether similar autonomous copies were present in other *D. melanogaster* genomes, we performed nucleotide BLAST searches against the NCBI whole-genome shotgun (wgs) contig database. These searches detected multiple hits corresponding to the degenerated chr3L-associated copies but also identified close matches in the sequenced genome of Bloomington Drosophila Stock Center stock #2969, which carries the classical *Bar*^1^ allele and is referred to hereafter as BL2969.

The BL2969 genome also showed a pronounced increase in the number of non-autonomous *Kodama* insertions, together with 9 autonomous copies located outside the chr3L locus containing the conserved degenerated copies. None of the nine *Nozomi* BL2969 copies occupied the same genomic positions as the 9 long A6 copies (Fig. 1E). Moreover, none of the *Nozomi* insertions in either A6 or BL2969 overlapped known *D. melanogaster* piRNA clusters (Fig. S2C).

A6 and BL2969 each contained more than 200 *Kodama* insertions and fragments, most of which fell within a narrow length range of 1420–1440 nt (Fig. 1F). Whereas dm6 contained only 13 *Kodama* insertions with both TIRs intact, A6 and BL2969 each contained more than 100 such insertions (Fig. 1F).

As in A6, the distribution of *Kodama* insertions relative to genes in BL2969 differed from that observed in dm6, with a larger fraction of insertions located near gene transcription start sites (TSSs) (Fig. S1C). Analysis of *Kodama* insertions relative to gene annotations across the DSPR founder genome panel further showed that, although only A6 displayed a pronounced expansion in *Kodama* copy number, several other genomes had insertion distributions more similar to that observed in A6 than to that observed in dm6 (Fig. S2D). This strongly suggests that *D. melanogaster* populations had experienced distinct histories of *Nozomi*-mediated *Kodama* mobilization and subsequent insertion retention.

Many expanded variants were highly similar to each other. In BL2969, one insertion variant was present at 32 different genomic loci without any nucleotide differences; when alignment gaps were disregarded, the same variant class encompassed 65 loci. Interestingly, despite their similar length and intact TIRs, most expanded *Kodama* variants in A6 and BL2969 did not show increased sequence similarity to each other. However, comparison of *Kodama* sequences from dm6 with expanded *Kodama* sequences from A6 and BL2969 indicated that the autonomous *Nozomi* mobilized pre-existing non-autonomous insertions as some of the most expanded insertions in BL2969 were similar to the insertions present in dm6 genome (Fig. S2E).

We did not detect any relationship between insertion sequence similarity and the genomic distance between insertions (Fig. S2F). This suggests that, unlike P elements, which preferentially transpose to nearby chromosomal sites, *Nozomi* does not show evidence of local transposition. Consistent with this interpretation, *Nozomi* insertions were distributed across all major chromosome arms, with a slight exception for BL2969, where no autonomous insertion was detected on chr2R (Fig. 1E).

We identified that there’s no strong hotspot adherence for *Nozomi/Kodama*, since only 7 out of 144 *Kodama* copies present in A6 genome share their location with any other genome from our combined panel.

### *Kodama* is defined by a precise central deletion

A defining feature of *D. melanogaster Kodama* elements, and a major rationale for recognizing them as a distinct mobile-element form, was that all short copies of approximately 1.4 kb identified across the 46 DSPR and DrosEU genomes carried the same precise internal deletion relative to the long autonomous variant, corresponding to the loss of a 1,380-bp central region (Fig. 2A, S3A).

**Figure 2.**
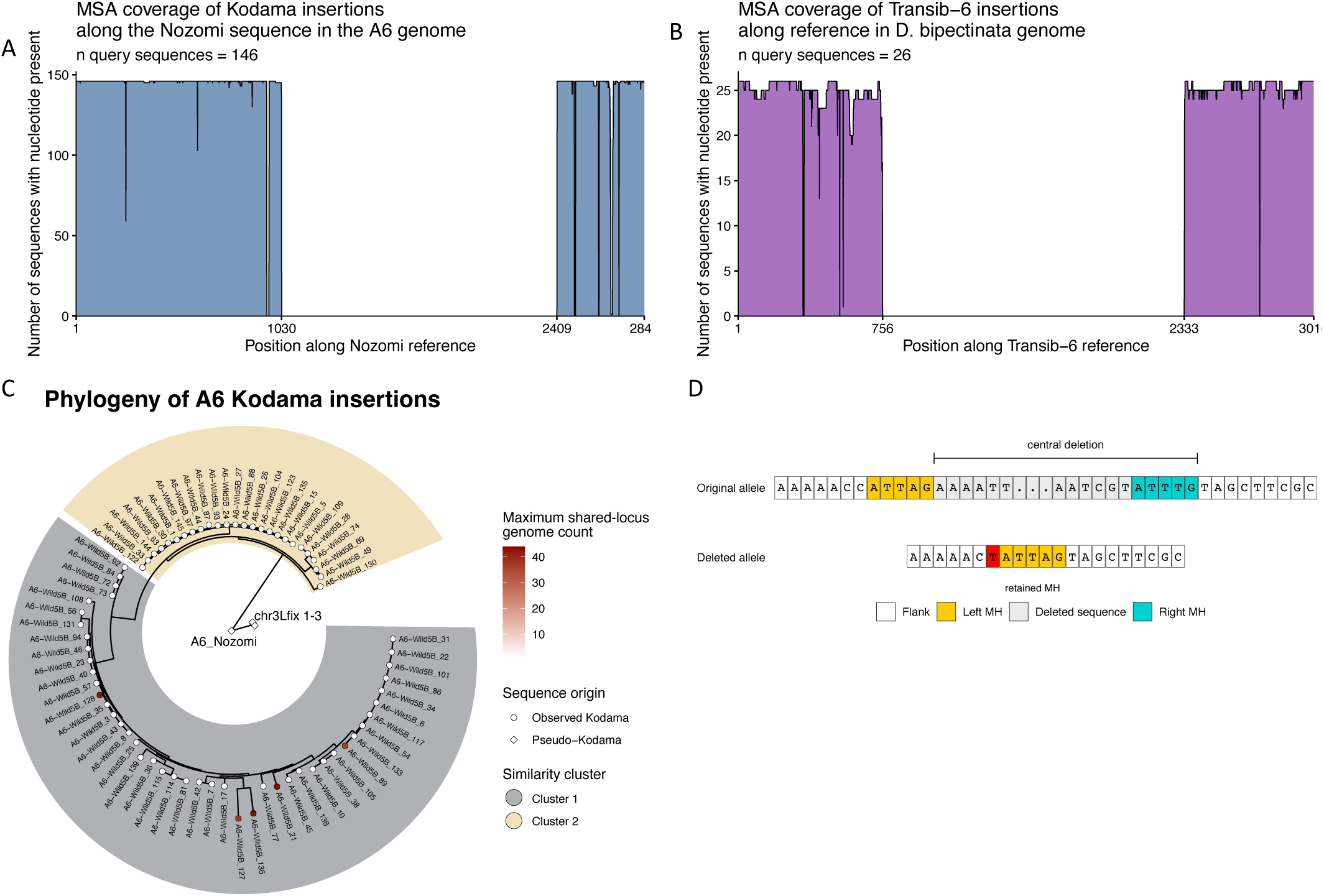
*Hopper/Kodama* elements in D. melanogaster show precise deletion of a central 1380 bp segment. (A) Multiple sequence alignment (MSA) coverage of all *Kodama* insertions bearing both TIRs in the *D. melanogaster* A6 genome along the autonomous *Nozomi* sequence. (B) MSA coverage of all *Transib-6* insertions bearing both TIRs in the *D. bipectinata* genome along the autonomous *Transib-6* sequence. (C) Phylogenetic relationships among unique short *Hopper* sequence variants in the A6 genome. The circular maximum-likelihood tree includes unique *Kodama* sequences together with pseudo- *Kodama* sequences generated by removing the central region from *Nozomi* insertions. Tip colour indicates the maximum insertion-locus prevalence associated with each sequence variant, defined as the largest number of genomes in which any corresponding insertion locus was detected. The two labelled similarity clusters were defined from pairwise K80 divergence among authentic *Kodama* sequences. (D) Microhomologies detected through comparison of *Nozomi* and *Kodama* sequences. Top: *Nozomi* sequence with central deletion and left and right microhomology sequences (MH) shown. Bottom: consensus sequence of *Kodama* shown with left MH retained. In red: T nucleotide showing a discrepancy with predicted MMEJ repair result.

To test whether this deleted form was also prevalent in broader population samples, we mapped DrosEU Pool-Seq genomic DNA reads from 48 pooled *D. melanogaster* populations to the autonomous *Nozomi* sequence. Read depth showed a clear depletion across the central deleted interval (Fig. S3B). In addition to the central deletion, the coverage profiles revealed several smaller depletion intervals, particularly in the 3′ region of the element, suggesting the presence of additional, less frequent internal deletions among natural *Kodama* copies. At the same time, the central deletion pattern inferred from short read coverage was smaller and, for unknown reasons, 5’ coverage of *Nozomi* was higher compared with its 3’ part.

A comparable central deletion pattern was observed in *Transib-6* from *Drosophila bipectinata*, the first *Nozomi* homolog identified in this study (Fig. 2B). The deleted interval in *Transib-6* was approximately 200 nt longer than that in *Nozomi*; however, because the full-length *Transib-6* element is also larger, the resulting non-autonomous derivatives were again concentrated around 1.4 kb (Fig. S3C, D). The independent enrichment of similarly sized non-autonomous derivatives in both lineages suggests that, in addition to full-length elements, efficient trans-mobilization may be restricted to a narrow secondary size range.

### Phylogenetic structure of *Kodama* insertions supports at least two amplification episodes

The widespread occurrence of *Kodama* copies carrying the same 1,380-bp central deletion raised the question of whether the current *Kodama* pool originated from a single expansion event or from multiple independent mobilization episodes. To address this, we extracted all unique *Kodama* insertion sequences from the A6 genome and reconstructed their phylogenetic relationships. We additionally generated four artificial deletion derivatives by removing the 1,380-bp central region from the autonomous *Nozomi* sequence and from the three deteriorated long copies on chromosome 3L. These sequences were included as reference points for assessing the relationship between the observed *Kodama* lineages and the known long *Nozomi* variants (Fig. 2C).

*Kodama* sequences separated into two well-defined clusters. Insertions detected at the same genomic positions across different genomes, representing shared insertion sites, in at least three of the 46 analyzed genomes were confined to Cluster 1, indicating that this cluster contains older, more broadly distributed insertions. By contrast, Cluster 2 was dominated by insertions restricted almost exclusively to the A6 genome, with only rare occurrences in other genomes. The distinct phylogenetic structure and cross-genome distribution of the two clusters therefore support at least two temporally separated episodes of *Kodama* mobilization.

The artificial deletion derivatives formed a separate, strongly supported group that was more closely related to Cluster 2 than to Cluster 1. Thus, the more recent *Kodama* cluster is genetically closer to the long *Nozomi* variants recovered in A6, whereas Cluster 1 represents a more diverged *Kodama* lineage. This topology alone cannot distinguish whether the 1,380-bp deletion arose once or independently on multiple occasions. Repeated mobilization of an existing deleted lineage and recurrent formation of similar deleted derivatives from related *Nozomi* variants are both compatible with the observed sequence relationships.

An additional unexpected pattern involved the three deteriorated long *Nozomi* copies on chromosome 3L. Long deteriorated copies were detected at the corresponding chr3L locus in at least 27 of the 46 analyzed genomes, yet their sequences remained relatively close to the newly identified autonomous *Nozomi* variants and substantially less diverged than the older *Kodama* cluster. Thus, sequence divergence of the long and short forms does not directly parallel their distribution across genomes, suggesting that the evolutionary histories of these two forms cannot be inferred from their present-day genomic distribution alone.

### Microhomology signatures suggest repair-mediated origin of *Kodama*

Inspection of the sequences flanking the recurrent internal deletion revealed 5 bp microhomologies immediately adjacent to the left and right deletion breakpoints (Fig. 2D). Similar breakpoint signatures were reported for the *Aedes aegypti* miniature inverted-repeat transposable elements (MITEs) *Gnome*, *Elf* and *Goblin*, which were inferred to originate from the autonomous DNA transposon *Ozma* through abortive gap repair followed by microhomology-mediated end joining (MMEJ)^35^. More broadly, studies in *Drosophila* have shown that Pol θ-dependent alternative end joining (alt-EJ)/MMEJ repairs double-strand breaks (DSBs) using short microhomologies and can generate templated junction products. Together, these observations suggest that *Kodama* most likely arose through an MMEJ- or synthesis-dependent MMEJ (SD-MMEJ)-like repair event acting on a full-length autonomous *Nozomi* copy. This interpretation is compatible with the broader ability of transposable elements (TEs) to exploit alt-EJ-related host repair pathways. In *Drosophila* oogenesis, long terminal repeat (LTR) retrotransposons use alt-EJ factors for DNA circularization and second-strand synthesis, showing that TE propagation can directly depend on this repair machinery^36^.

Notably, most *Kodama* copies also carry a T, rather than the C present in the *Nozomi* copies recovered from A6 and BL2969, immediately upstream of the retained left microhomology (Fig. 2D, S4A, B). This suggests that the amplified short lineage likely arose from a closely related but non-identical ancestral autonomous copy, or from an early deleted derivative that later expanded. Additional single-nucleotide variants shared among *Kodama* copies but absent from *Nozomi* further support this interpretation (Fig. S4A, B). Of note, similar microhomology signatures couldn’t be found for *Transib-6* in *D. bipectinata*. One possible explanation could come from the fact that the short *Transib-6* insertions are more diverged from the identified autonomous insertion and overall higher accumulation of shorter degraded insertions in *D. bipectinata* genome (Fig. S3C) which hints at the possibility that the autonomous insertion found by us is quite diverged from the ancestral autonomous transposon.

The predominance of 1.4 kb non-autonomous *Kodama* insertions in *D. melanogaster*, together with the enrichment of similarly sized short derivatives in *D. bipectinata Transib-6* (Fig. S3D), suggests that internally deleted derivatives can become the major mobilized form of this *Transib* lineage. These elements retain terminal inverted repeats (TIRs) but lack transposase-coding capacity, resembling MITEs in their dependence on an autonomous transposase provided in trans. However, their length exceeds the size range typically associated with canonical MITEs. Thus, short copies are best interpreted as unusually long MITE-like derivatives generated by internal deletion of an autonomous *Transib* element.

Together, these results support a model in which a repair-mediated deletion generated a non-autonomous *Kodama* derivative that retained the terminal and internal sequences required for mobilization and subsequently became a predominant mobilized form in some *D. melanogaster* genomes.

### The predicted *Nozomi* transposome structure closely resembles *HzTransib* crystal strand-transfer complex

We used AlphaFold 3 (AF3) to model the *Nozomi* transposome as a transposase homodimer bound to two double-stranded TIRs and four Mg²⁺ ions (Fig. 3A). Despite only 42.8% amino- acid identity and the substantial phylogenetic separation between *Nozomi* and *HzTransib*, the well-characterized *Transib* DNA TE found in the corn earworm *Helicoverpa zea*, the predicted *Nozomi* complex closely matched the experimentally determined *HzTransib* strand-transfer complex (PDB 6PR5^17^). Structural superposition aligned 443 Cα atoms with an RMSD of 1.54 Å (Fig. 3B). Moreover, after fitting only the first protein subunit, the second *Nozomi* subunit overlapped the corresponding *HzTransib* subunit with an RMSD of 1.52 Å over 432 Cα atoms, indicating strong conservation of the dimer architecture (Fig. 3B).

**Figure 3.**
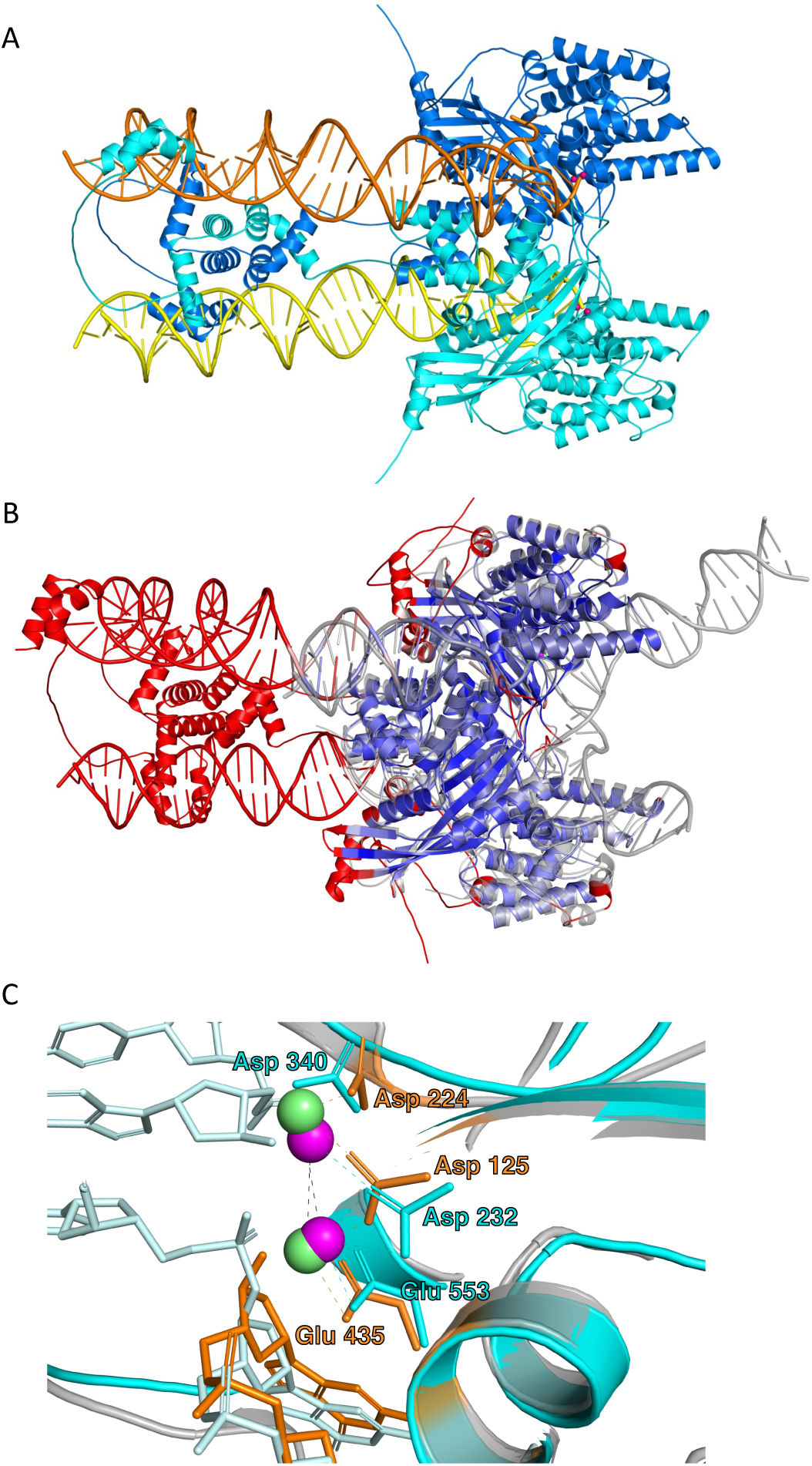
Predicted *Nozomi* transposase structure closely resembles the solved *HzTransib* structure. (A) AlphaFold3 predicted structure of *Nozomi* transposase dimer together with two TIRs and four Mg²⁺ ions. (B) Structural superposition of the template-free AF3 *Nozomi* model with the experimentally determined *HzTransib* strand-transfer complex (PDB 6PR5). The *Nozomi* transposase dimer and bound DNA are coloured according to their local geometric displacement from 6PR5, with blue indicating the closest agreement, white intermediate displacement, and red the most divergent regions. The 6PR5 protein–DNA complex is shown in semi-transparent grey. (C) Structural superposition of catalytically active centers of AF3 predicted structure of *Nozomi* transposase and *HzTransib* strand-transfer complex (PDB 6PR5). *Nozomi* is colored in cyan, *HzTransib* is colored in orange.

The active-site geometry was even more highly conserved. The predicted *Nozomi* catalytic residues D232, D340, and E553 coincided with *HzTransib* D125, D224, and E435, respectively, with a combined Cα RMSD of 0.23 Å (Fig. 3C). The two Mg²⁺ ions predicted in each *Nozomi* active site were positioned 0.98 and 0.59 Å from the corresponding experimental ions in 6PR5. Thus, the *Nozomi* model reproduced both the catalytic DDE geometry and the two-metal centre characteristic of the assembled strand-transfer complex.

The initial AF3 prediction used the apo *HzTransib* homodimer structure (PDB 6PQN^17^) as an inference-time protein template. To test whether the close correspondence to 6PR5 depended on this template, we repeated the prediction with structural templates disabled while retaining the same molecular input and random seed. The template-free model was essentially unchanged and superimposed onto 6PR5 with a Cα RMSD of 1.53 Å over 443 residues. Its D232/D340/E553 catalytic triad matched the corresponding *HzTransib* residues with an RMSD of 0.23 Å, while the two predicted Mg²⁺ ions were positioned 0.95 and 0.56 Å from their experimental counterparts. The predicted *Nozomi* complex therefore robustly adopts a strand-transfer-like *Transib* conformation regardless of whether the apo *HzTransib* structure is supplied as an inference-time template.

### Comparative searches identify *Nozomi*-related *Transib* elements in dipterans

We included all Drosophilidae (phylogeny in Fig. 4A) in nucleotide BLAST searches against the NCBI whole-genome shotgun contig database using *Nozomi* sequence as the query, and identified only a single close hit outside *D. melanogaster*, corresponding to the R32 strain of *Drosophila mauritiana* (94.6% AA identity) (Fig. 4B). The presence of only a single detectable copy may indicate that *Nozomi* failed to expand after its introduction, potentially because it inserted into a piRNA-producing locus or another repressive chromatin environment.

**Figure 4.**
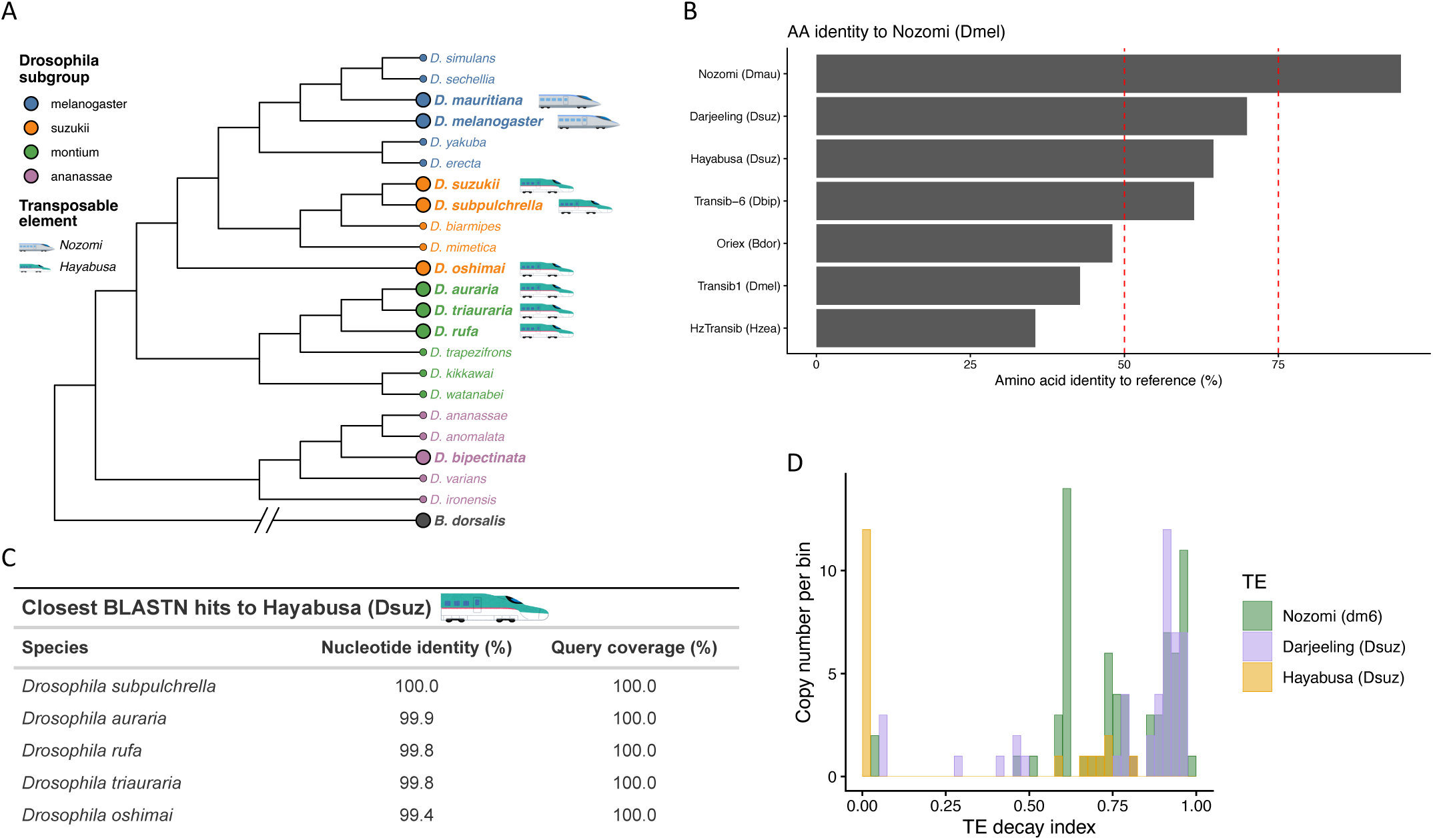
*Nozomi* homologs in other dipteran species. (A) Phylogeny of the dipteran species examined in this study. The cladogram was obtained by pruning the phylogenomic ASTRAL tree of Kim et al.^68^ to representative species from the melanogaster, suzukii, montium, and ananassae subgroups. Species included in the analyses are highlighted by larger tip symbols and bold labels. *Bactrocera dorsalis*, a member of Tephritidae family, shown as the outgroup in gray. Train icons indicate species in which *Nozomi* or *Hayabusa* was identified. Train illustrations adapted from Irasutoya (いらすとや) and used in accordance with its terms of use. (B) Amino acid identity of *Nozomi* homologs identified in *D. mauritiana, D. suzukii*, *D. bipectinata* and *B. dorsalis*. (C) Table with the closest nucleotide BLAST hits to the *D. suzukii Hayabusa* DNA transposon across species of the *Drosophila* montium and suzukii subgroups. (D) Distribution of TE decay metric for *Hayabusa* and *Darjeeling* copies in the *D. suzukii* genome and *Nozomi* copies in the *D. melanogaster* dm6 genome.

Using protein BLAST and translated BLAST searches with the *Nozomi* transposase amino acid sequence, we identified homologous *Transib*-family elements in three additional dipteran species: two elements in *Drosophila suzukii*, with amino acid similarities of 82.3% and 72.9%, and one element in *Bactrocera dorsalis*, with amino acid similarity of 62.7% (Fig. 4B). We named the *Transib*-superfamily homolog identified in *B. dorsalis Oriental Express,* or *Oriex*, as it continues with railway analogy and *B. dorsalis* is also known as oriental fruit fly.

In *D. suzukii*, we named the closer *Nozomi* homolog *Darjeeling* and the more distant homolog *Hayabusa*. *Darjeeling* likely represents an ancient invasion, as we detected few intact autonomous copies in the *D. suzukii* genome, while *Hayabusa* is most likely a novel TE family introduced in *D. suzukii* genome recently.

Multiple sequence alignment of *Nozomi* homologs, together with *D. melanogaster Transib1* and *Hztransib*, revealed a conserved DDE transposase catalytic core, consistent with the two-metal-ion mechanism characteristic of cut-and-paste DNA transposases (Fig. S5A). The amino acid similarity between the *Nozomi* homologs increased even more when we compared only the conserved RNAse H-like fold domains of the transposases containing first two key residues of the catalytic triad (Fig. S5B). All identified *Nozomi* homologs also contained N-terminal helical domains absent from *Transib1* and *Hztransib*.

As with *D. melanogaster Nozomi*, we could not find any domain matches in Interproscan for all identified homologs. However, running HHPRED produced 100% probability matches with *Hztransib,* RAG1L DNA transposase from *Branchiostoma belcheri* and RAG1(V(D)J recombination-activating protein 1), from *Mus musculus* (Supp. Table 1).

### *Hayabusa* shows signatures of recent horizontal transfer across suzukii and montium subgroups

Nucleotide BLAST searches detected near identical *Hayabusa* sequences across the *Drosophila* suzukii and montium species groups (Figure 4C). These copies showed minimal nucleotide divergence despite their distribution across multiple species, a pattern consistent with recent horizontal transfer (HT) rather than long-term vertical inheritance. When genomic DNA reads from the *D. suzukii* population-structure dataset were mapped to the autonomous *Hayabusa* sequence, coverage remained comparatively uniform across the element, unlike *Nozomi* in *D. melanogaster* (Fig. S5C, compare with Fig. S3A).

To further compare the recent activity and structural preservation of *Hayabusa* with related elements, we analyzed insertion divergence and decay for three *Transib*-like families: *Nozomi/Kodama* in *D. melanogaster* and the two closest *D. suzukii* homologs, *Darjeeling* and *Hayabusa*. First, we calculated Kimura two-parameter (K80) distances between individual copies and their corresponding consensus sequences and plotted copy numbers in 0.005 K80 bins (Fig. S5D). This provided a measure of nucleotide divergence among recovered copies. Comparison of *Nozomi* K80 distributions among dm6, A6, and BL2969 genomes revealed the expected increase of intermediately diverged *Kodama* copies in A6 and BL2969 (Fig. S5E).

Because nucleotide divergence alone doesn’t capture large deletions or truncated BLAST matches, we also calculated a TE decay index combining sequence divergence and copy completeness. In this metric, an intact full-length insertion identical to the consensus has a decay value of 0, whereas increasing K80 distance and decreasing alignment length increase the score. Thus, TE decay is not intended as an absolute molecular clock, but as a relative measure of insertion deterioration that jointly reflects nucleotide divergence and structural completeness.

The three elements showed clearly distinct decay profiles. *Hayabusa* insertions in *D. suzukii* were concentrated close to zero in both K80 distance and TE decay (Fig. 4D, S5D), indicating the presence of largely full-length, near identical copies. This pattern is consistent with recent or ongoing activity of *Hayabusa* in the analyzed genome. In contrast, *Darjeeling* in *D. suzukii* and *Nozomi* in *D. melanogaster* (dm6 genome) showed broader and more shifted decay distributions, with few copies occupying near zero decay class (Fig. 4D). These profiles indicate that most recovered *Darjeeling* and *Nozomi/Kodama* insertions are either diverged from their autonomous counterparts, structurally incomplete or both.

Together, the broad distribution of minimally diverged *Hayabusa* copies across suzukii and montium subgroups and the low-decay profile in *D. suzukii* support a recent spread of this element, most likely through horizontal transfer. By contrast, *Darjeeling* and *Nozomi* are dominated by older or structurally deteriorated copies in the analyzed reference genomes.

### Target site recognition varies across *Nozomi*-related *Transib* elements

Despite substantial amino acid similarity among the predicted transposases, *Nozomi* homologs differed unexpectedly in their target-site duplication (TSD) length and sequence preference (Table S2). Canonical *Transib* elements have been distinguished from the related CACTA– Mirage–Chapaev (CMC) group in part by their 5 bp TSDs, whereas CMC elements show a broader 2–4 bp TSD range^37^. Consistent with this canonical *Transib* pattern, *D. melanogaster Nozomi* is associated with a 5 bp CARTG TSD. In contrast, the *D. suzukii* element *Hayabusa* and *D. bipectinata Transib-6* showed 3 bp TSD consensus motifs, ANT and AST, respectively, whereas *Oriex* from *B. dorsalis* was associated with an apparent TA dinucleotide TSD, a target-site signature more typical of Tc1/mariner elements than of *Transib*^38^ (Table S2).

These observations suggest that target-site recognition may be more variable within the *Hopper*- related *Transib* lineage than expected from previously described *Transib* elements. In particular, the 3 bp TSDs of *Hayabusa* and *Transib-6* overlap the length range reported for CMC elements, while the apparent TA TSD of *Oriex* represents an even stronger departure from the canonical 5 bp *Transib* pattern. Notably, the *Nozomi* homologs with non-canonical TSDs also had terminal inverted repeats (TIRs) beginning with G rather than the C nucleotide typical of described *Transib* termini (Table S2). These observations indicate that target-site recognition is extremely variable within the *Nozomi*-related *Transib* lineage. The 3-bp TSDs of *Hayabusa* and *Transib-6* and the apparent TA dinucleotide TSD of *Oriex* depart from the canonical 5-bp pattern, despite clear *Transib* affinity at the transposase level. The accompanying variation in TIR termini further suggests that both terminal architecture and target-site selection have diversified within this lineage.

## Discussion

More than three decades after the initial description of *Drosophila melanogaster Hopper* as a short non-autonomous DNA transposon^28^, we identified the corresponding full-length autonomous element and resolved the relationship between the two forms. We designate the autonomous element *Nozomi* and the previously described short *Hopper* form *Kodama*. *Nozomi* encodes a 648-amino-acid DDE transposase and is firmly placed within the *Transib* superfamily by protein-sequence similarity, phylogenetic position and the close structural correspondence of its predicted transposome to the experimentally determined *Helicoverpa zea HzTransib* strand-transfer complex. The genomic histories of the autonomous and non-autonomous forms, however, differ. While all the analyzed *D. melanogaster* genomes contain at least some number of *Kodamas*, autonomous *Nozomi* is restricted to a small number of *D. melanogaster* genomes, where its presence in A6 and BL2969 is accompanied by extensive and independent amplification of *Kodama*. All examined *Kodama* copies share the same precise 1,380-bp central deletion, their phylogenetic structure in A6 genome records at least two episodes of mobilization, and microhomology at the deletion boundaries suggests a repair-mediated origin involving alternative end joining. A related *Transib-6* lineage in *Drosophila bipectinata* independently shows enrichment for a similarly sized non-autonomous derivative, suggesting that mobilization of shortened elements may be a recurring feature of this lineage. At the same time, the related *Hayabusa* element presents a strikingly different evolutionary outcome: minimal sequence decay with no apparent centrally deleted copies and broad distribution across closely related *Drosophila* species are consistent with a recent and highly successful horizontal invasion. Together, these findings define a previously unrecognized *Transib* lineage in which closely related elements show markedly different invasion success while repeatedly giving rise to abundant shortened non-autonomous derivatives.

### *Nozomi* provides an experimentally tractable model for *Transib*-RAG evolution

The evolutionary relationship between *Transib* transposases and RAG1 makes *Nozomi* of particular interest beyond its recent activity in *Drosophila*. Despite substantial sequence divergence, the predicted *Nozomi* transposome closely reproduces the catalytic architecture and dimer organization of the experimentally determined *Hztransib* strand-transfer complex. At the same time, the *Nozomi* transposase retains considerable sequence similarity to vertebrate RAG1, including 30.4% amino acid similarity to mouse RAG1. The availability of active autonomous copies, genetically tractable *D. melanogaster* strains and extensive molecular tools therefore makes *Nozomi* a potentially useful experimental system for investigating conserved features of *Transib* transposition and their relationship to the evolution of RAG-mediated recombination.

### Recent reintroduction or reactivation of an ancestral copy?

An important question requiring addressing is whether *Nozomi* was reintroduced into *D. melanogaster* by horizontal transfer or reactivated from ancestral genomic copies. Several observations favor recent reintroduction. Highly similar *Nozomi* sequences occur in distinct *D. melanogaster* strains and in *D. mauritiana*, whereas the *Kodama* copies shared across many *D. melanogaster* genomes are substantially more diverged. Moreover, no *Kodama* copy was identical to the autonomous variants recovered from A6 and BL2969, and the active long copies in these two genomes did not share insertion positions. Together, these findings argue against expansion from a single ancestral genomic copy and instead suggest that *Nozomi* entered A6 and BL2969 through separate events, most plausibly through independent horizontal transfers of closely related variants.

The coexistence of divergent ancestral copies and closely related autonomous *Nozomi* variants raises the possibility that this lineage has entered Drosophila melanogaster more than once. A comparison with recent TE invasions is informative. *Transib1* spread rapidly through natural D. melanogaster populations following its introduction^21^, whereas autonomous *Nozomi* remains restricted to a small number of strains. *Tirant* provides a closer precedent for repeated invasion: degraded copies from an ancient invasion persisted in D. melanogaster and produced piRNAs, yet they did not prevent the spread of a distinct canonical *Tirant* variant during the past century^39^. Similarly, piRNAs derived from ancestral, diverged *Nozomi* copies may provide incomplete protection against an incoming autonomous variant. The presence of old genomic remnants may therefore be compatible with, rather than protective against, recurrent horizontal introduction of a related TE lineage.

### Why did *Nozomi* fail to spread broadly?

The restricted distribution of autonomous *Nozomi* suggests that its recent introductions did not result in a species-wide invasion. Several non-exclusive explanations are possible. The autonomous variants may have entered populations with limited opportunity for further spread, may have imposed sufficient fitness costs to be rapidly eliminated, or may have been suppressed soon after introduction by the piRNA pathway. The extensive amplification of short non-autonomous copies may also have contributed to this outcome. Because these deleted variants retain the terminal sequences required for mobilization but do not encode transposase, they may compete with autonomous elements for the same transposition machinery. Their expansion could therefore limit amplification of the autonomous form while simultaneously increasing the genomic burden of *Hopper*-derived insertions.

The prevalence of the same approximately 1.4 kb deleted architecture across *D. melanogaster* populations as well as for *Transib-6* in *D. bipectinata* supports the idea that this non-autonomous form became the principal mobilized substrate of the *Nozomi* transposase. In this respect, short *Kodama* elements resemble unusually long MITE-like derivatives: they lack protein-coding capacity, depend on an autonomous transposase supplied in trans and can reach high genomic copy numbers. Their predominance suggests that the evolutionary outcome of autonomous mobile element introduction is determined not only by the interaction between the transposon and host defense, but also by competition between autonomous and non-autonomous members of the same TE family.

### Did *Kodama* arise once or repeatedly?

The evolutionary history of the central *Nozomi* deletion that produced *Kodama*s remains difficult to resolve. Phylogenetic analysis of *Kodama* insertions suggests several distinct episodes of amplification, but these bursts do not necessarily imply that the 1,380 bp deletion arose independently during each episode. They could instead reflect repeated mobilization of an already existing non-autonomous lineage following separate introductions or reactivations of *Nozomi* transposase.

Several observations favor a single ancestral origin of the deleted variant. All short *Hopper* copies examined carried the same deletion boundaries, and we detected no alternative long-to-short intermediates that would indicate repeated formation of similar deletions from different autonomous copies. Moreover, apart from the degenerated long copies at the chr3L locus, which have lost the intact 5′ terminal inverted repeat required for transposition, the analyzed genomes contained no widespread diversity of ancestral long *Nozomi* variants from which the short copies could have arisen independently. Instead, autonomous copies were restricted to a small number of genomes, whereas the centrally deleted form was broadly distributed and substantially more abundant.

Under the simplest model, the central deletion therefore arose once in an ancestral autonomous *Nozomi* copy, producing a non-autonomous derivative that persisted in *D. melanogaster* populations after functional autonomous copies were lost or silenced. Subsequent introductions of related autonomous *Nozomi* variants could then have remobilized this pre-existing short lineage, generating the distinct amplification bursts observed in the phylogeny. This model would also explain why the expanding short copies are not identical to the autonomous variants currently present in A6 and BL2969: the deleted lineage may predate these autonomous copies and have been mobilized repeatedly in trans.

Nevertheless, recurrent generation of the same deletion cannot be completely excluded. The presence of short microhomologies at the deletion boundaries may strongly favor repair at these particular sites, potentially allowing similar deletions to arise independently after multiple *Nozomi* introductions. Distinguishing between these models will require reconstruction of the relationships among deletion-flanking haplotypes, broader sampling of autonomous *Nozomi* variants, and identification of any rare intermediates linking full-length and short copies. At present, however, the shared deletion boundaries, broad distribution of short copies and scarcity of diverse long ancestral variants are most consistent with a single origin of the deleted lineage followed by repeated episodes of trans-mobilization.

### TIR and TSD architectures are more evolutionarily labile than the *Transib* catalytic core

Cut-and-paste DNA transposons are classified primarily from transposase sequence relationships, while TSD length and the terminal nucleotides of their TIRs are commonly used as supporting diagnostic features. Early studies defined *Transib* elements by their characteristic 5- bp TSDs, frequently associated with GC-rich target sequences^15^, and subsequently identified a conserved RSS-like TIR architecture containing a CACAATG-like terminal heptamer^13^.

However, the *Nozomi*-related elements characterized here show that neither feature is invariant within the broader *Transib* lineage. *Nozomi* was associated with a 5-bp CARTG TSD, whereas *Hayabusa* and *Transib-6* were associated with 3-bp ANT and AST motifs, respectively, and *Oriex* generated an apparent TA dinucleotide duplication more commonly associated with Tc1/mariner elements (Table S2). Their TIR sequences were likewise considerably more variable than expected from previously characterized *Transib* families. Despite this variation, the corresponding transposases consistently grouped with *Transib* proteins, and the predicted *Nozomi* catalytic centre closely superimposed on that of *HzTransib* despite only 42.8% amino acid identity. This contrast suggests that the catalytic chemistry of transposition is more strongly constrained than the molecular determinants of transposon-end recognition and target capture.

TSD length reflects the spacing between the two strand-transfer reactions within the assembled transpososome, whereas TIR recognition and target-site selection depend on additional DNA-contacting surfaces. Structural analysis of *HzTransib* showed that several non-catalytic regions, including the C-terminal tail, ZnB domain and α9–α10 loop, contact the TIR or target DNA and contribute to substrate positioning^17^. The notable divergence of non-catalytic regions among *Nozomi*-related transposases, including their N-terminal helical domains, could therefore permit changes in terminal-sequence recognition and integration-site geometry without disrupting the conserved DDE catalytic core, although the contribution of these N-terminal domains remains to be tested. *Oriex* provides a particularly clear example: its apparent TA duplication resembles the canonical insertion footprint of Tc1/mariner elements, yet its transposase sequence and phylogenetic placement support *Transib* ancestry. TSD length and TIR sequence should therefore be regarded as evolutionarily variable mechanistic traits that support, but should not override, classification based on transposase homology and catalytic architecture.

## Materials and Methods

### Identification of transposable element insertions

Transposable element insertions were identified in assembled *Drosophila melanogaster* genomes using Earl Grey v.7.1.0^40^. The analyzed assemblies included founder strains from the Drosophila Synthetic Population Resource^29,30,41^ (DSPR; NCBI BioProject PRJNA418342) and population-scale long-read genome assemblies of natural *D. melanogaster* strains collected predominantly across Europe^27^ (NCBI BioProject PRJNA559813). The DSPR BioProject contains genome assemblies for 14 founder strains, whereas PRJNA559813 contains the scaffolded assemblies and sequencing data for the 32-strain long-read panel.

Each genome assembly was analyzed independently using 16 computational threads. A custom *D. melanogaster* TE consensus library containing the TE families examined in this study was supplied to Earl Grey as the initial library of known elements using the `-l` option. Potentially spurious annotations shorter than 100 bp were removed using the `-m yes` option. Generation of a soft-masked genome was disabled (`-d no`), and the optional HELIANO module for Helitron detection was not run (`-e no`). De novo consensus-sequence clustering was not enabled, and all other Earl Grey parameters were left at their default values. Final filtered and merged Earl Grey annotations were used as the set of TE insertion coordinates for downstream analyses.

### Coordinate liftover and genomic feature annotation of TE insertions

To compare TE insertion positions among independently assembled *Drosophila melanogaster* genomes, insertion coordinates were converted to the dm6 reference coordinate system. Each source genome was aligned to the dm6 assembly using minimap2^42^ with the `asm20` assembly-to-assembly alignment preset and base-level alignment information retained using the `--cs` option. The resulting pairwise alignments were stored in PAF format. TE coordinates in BED format were subsequently transferred from the source assembly to dm6 using `paftools.js liftover`, with a minimum lifted interval length of 20 bp. A custom wrapper retained the original BED metadata after coordinate conversion, recorded intervals that could not be mapped, and sorted successfully lifted intervals by genomic position.

For analysis of genomic feature distribution, lifted *Kodama* insertion loci were represented by their midpoint coordinates and reduced to 1-bp intervals, thereby assigning each insertion to a genomic feature according to its insertion site rather than the complete TE span. Only standard dm6 chromosomes were retained, and chromosome naming was converted to the UCSC convention. Insertion sites were annotated using ChIPseeker^43^ against the *D. melanogaster* dm6 Ensembl gene annotation provided by `TxDb.Dmelanogaster.UCSC.dm6.ensGene`, with gene identifiers annotated using `org.Dm.eg.db`. Promoters were defined as the region extending 1 kb upstream and downstream of the transcription start site. Insertions were classified by ChIPseeker into promoter, exon, intron, UTR, downstream, and distal intergenic categories, and the relative distribution of these feature classes was compared among DSPR founder genomes and the dm6 reference.

### Whole-genome synteny analysis

Whole-genome synteny between the A6-Wild5B, A1 (Canton-S), and dm6 assemblies was assessed using minimap2^42^ and SyRI^44^. Assembly-to-assembly alignments were generated with the `asm5` preset, and structural rearrangements were identified from sorted BAM alignments using SyRI. The resulting synteny information was inspected to characterize the genomic context of a large inversion involving the A6 genome.

### Phylogenetic analysis of *CMC–Transib* DDE transposases

Protein sequences representing *CMC–Transib* DDE DNA transposases were obtained from the RepeatPeps protein library distributed with RepeatMasker^45^. Sequences annotated as DNA/CMC-Chapaev, DNA/CMC-Chapaev-3, DNA/CMC-EnSpm, DNA/CMC-Mirage, or DNA/CMC-Transib were extracted from the library. Additional curated protein sequences, including the *Drosophila melanogaster Nozomi* transposase, selected *Transib* transposases, and RAG1 proteins, were added from a separate FASTA file. Sequence identifiers were standardized, and noncanonical amino acid characters and terminal stop symbols were converted to `X`. Protein sequences were aligned with MAFFT^46^ (v7.526) using the L-INS-i strategy, with local pairwise alignment information and 1,000 iterative refinement cycles (`--localpair -- maxiterate 1000`). Poorly aligned positions were removed using ClipKIT (v2.12.2) in `smart- gap` mode, which determines the gap-frequency threshold dynamically while retaining phylogenetically informative alignment positions^47^.

A maximum-likelihood phylogeny was inferred from the trimmed amino acid alignment using IQ-TREE 3^48^. The analysis used the fixed LG+F+I+G4 substitution model, comprising the LG amino acid replacement matrix^49^, empirical amino acid frequencies, a proportion of invariant sites, and gamma-distributed rate heterogeneity approximated using four discrete categories. Branch support was assessed using 1,000 ultrafast bootstrap replicates^50^ and 1,000 SH-like approximate likelihood-ratio test replicates^51^. The random-number seed was fixed at 1 to ensure reproducibility.

The resulting consensus tree was imported into R using the ape package^52^. Any root present in the imported tree was removed, and the phylogeny was visualized as an unrooted cladogram with branch lengths suppressed using ggtree^53^. *Nozomi*, selected *Transib* transposases, and RAG1 proteins were highlighted for visualization.

### Phylogenetic analysis of *Kodama* insertions

Phylogenetic relationships among *Kodama* insertions in the A6 genome were examined using unique nucleotide sequences. Identical short-copy sequences were collapsed, retaining one representative sequence for phylogenetic analysis. Prior to multiple sequence alignment, short sequences were oriented in the same direction as the autonomous *Nozomi* element.

To assess the relationship between *Nozomi* and *Kodama* variants, pseudo-short sequences were generated from the autonomous A6 *Nozomi* insertion and three deteriorated long insertions located on chromosome 3L. For each long insertion, the central region corresponding to the characteristic deletion in *Kodama* was removed, and the retained 5′ and 3′ regions were concatenated. These pseudo-*Kodama* sequences were included as phylogenetic references but were excluded from insertion-locus prevalence calculations.

Sequences were aligned with MAFFT^46^ using the automatic strategy selection option (`--auto`). A maximum-likelihood phylogeny was inferred from the resulting alignment using IQ-TREE 3^54^ under the GTR+G nucleotide substitution model. Branch support was evaluated using 1,000 ultrafast bootstrap replicates^50^ and 1,000 SH-like approximate likelihood-ratio test replicates^51^. The consensus tree was visualized as an unrooted circular phylogeny using the `ggtree`^53^ package in R.

Pairwise sequence divergence was calculated from the MAFFT alignment under the Kimura two-parameter model^55^ using the `dist.dna` function from the `ape`^52^ R package. Sites containing gaps or missing data were not removed on a pairwise basis. Average-linkage hierarchical clustering of the K80 distance matrix was used to identify the principal sequence clusters. The pseudo- *Kodama* sequences derived from *Nozomi* insertions formed a separate cluster and were therefore not included when assigning the two principal clusters of naturally occurring *Kodama* insertions. Each natural *Kodama* sequence was associated with its genomic insertion-locus group and with the number of surveyed genomes containing an insertion at that locus. When the same nucleotide sequence occurred at more than one insertion-locus group, the highest locus prevalence among the corresponding groups was assigned to the representative sequence. This maximum insertion- locus prevalence was used to annotate tips in the phylogenetic tree. Sequence manipulation and downstream analyses were performed in R using the Biostrings^56^ and dplyr^57^ packages.

### Population sequencing coverage analysis

Population-level coverage of *Nozomi* in *D. melanogaster* and *Hayabusa* in *D. suzukii* was assessed using publicly available whole-genome sequencing datasets. For Nozomi, sequencing reads from DrosEU populations^58^ were analyzed, whereas Hayabusa coverage was assessed across population genomic datasets of *D. suzukii*^59^. Only R1 reads from paired-end libraries were used. Reads were quality-filtered with fastp^60^ using a minimum qualified-base Phred score of 10 and mapped to the corresponding full-length TE reference sequence with Bowtie2 in `--very- sensitive` mode^61^. No mapping-quality filtering was applied.

To account for differences in sequencing depth among samples, the same reads were independently mapped to the corresponding host reference genome, dm6 for *D. melanogaster* and the *D. suzukii* reference genome for *Hayabusa*. Per-base TE coverage was calculated using `samtools depth`^62^ and normalized per million reads mapped to the host genome. For comparison of coverage profiles among samples, normalized coverage at each position was subsequently divided by the maximum coverage within that sample. For the *D. suzukii* analysis, samples covering more than 95% of the *Hayabusa* reference sequence were retained for the combined coverage visualization.

### AlphaFold 3 modelling of the *Nozomi* transposome

The *Nozomi* transposome was modelled using AlphaFold 3 (AF3) through AlphaFold Server. The input comprised two identical *Nozomi* transposase chains, two copies of each complementary 37-nt terminal inverted repeat (TIR) strand, corresponding to two double-stranded TIR molecules, and four Mg²⁺ ions. AF3 jointly predicts complexes containing proteins, nucleic acids and ions, allowing the complete protein–DNA–metal complex to be modelled in a single calculation^63^. The initial prediction was generated using the random seed 1276554233 with structural-template searching enabled. Inspection of the downloaded template files showed that the apo *HzTransib* homodimer structure (PDB 6PQN) had been selected as the highest-ranking protein template.

To test whether the predicted conformation depended on the *HzTransib* template, the calculation was repeated with an otherwise identical AlphaFold Server JSON input in which `useStructureTemplate` was set to `false` for the *Nozomi* protein chains. The protein and DNA sequences, molecular stoichiometry, Mg²⁺ content and random seed were retained unchanged. The absence of structural templates in the resulting prediction was confirmed by inspection of the downloaded output directory. The highest-ranked model from each run was used for subsequent analyses.

### Structural comparison with the *HzTransib* strand-transfer complex

Predicted *Nozomi* complexes were compared with the experimentally determined *HzTransib* strand-transfer complex (PDB 6PR5), which contains a transposase dimer, transposon-end and target DNA, and two catalytic Mg²⁺ ions per active site. Structural analyses were performed in PyMOL using custom Python scripts. *Nozomi* chain A was superimposed onto 6PR5 chain A using corresponding protein Cα atoms. The principal global RMSD was calculated without iterative outlier removal (`cycles = 0`) to retain all residue pairs identified by the structural alignment. A refined conserved-core RMSD was additionally calculated using a 2 Å rejection cutoff and five refinement cycles. Because PyMOL refinement removes poorly matching atom pairs, the unpruned RMSD was used as the primary measure of global structural similarity. Conservation of the dimer architecture was assessed without performing an independent fit of the second subunit. After superimposing *Nozomi* chain A onto 6PR5 chain A, the Cα RMSD between *Nozomi* chain B and 6PR5 chain E was calculated in the resulting coordinate frame with structural transformation disabled. Thus, agreement of the second subunit reflected conservation of its position relative to the fitted subunit rather than a separate optimization.

For residue-level analysis, the distance between each pair of corresponding Cα atoms in the unpruned structural alignment was calculated. Alignment coverage was expressed relative to the total number of Cα atoms in each protein chain, and the proportions of aligned residues separated by ≤1, ≤2, ≤3 and ≤5 Å were determined. Continuous well-superimposed regions were defined as runs of at least eight sequential residue pairs with Cα displacement ≤2 Å. Residue-wise displacement values were stored in the B-factor field of a working copy of the predicted structure and used to colour the complex from blue for close agreement through white to red for increasing displacement.

The catalytic centres were evaluated using the established *HzTransib* catalytic residues D125, D224 and E435, which mapped to *Nozomi* D232, D340 and E553, respectively. Their combined Cα RMSD was calculated in the global protein-alignment frame without further fitting. To assess alignment-independent catalytic geometry, pairwise Cα distances among the three catalytic residues were calculated in both structures. Metal coordination was assessed by measuring the shortest distance between the carboxylate oxygen atoms of each catalytic residue and the Mg²⁺ ions associated with the corresponding active site.

Mg²⁺ ions were not included in the protein superposition. Following protein-only alignment, the two predicted ions in each *Nozomi* active site were paired with the two experimental ions in 6PR5 using the one-to-one assignment that minimized the sum of Mg²⁺–Mg²⁺ distances. The RMSD of the resulting two-ion assignment was then calculated, providing an independent measure of predicted metal placement.

A separate local comparison of the catalytic domain was performed using all structurally mapped Cα pairs whose corresponding 6PR5 residues lay within 18 Å of either experimental Mg²⁺ ion, together with the three catalytic residues. These explicitly paired coordinates were fitted using a rigid-body Kabsch superposition without outlier removal. Catalytic-domain RMSD, catalytic-triad RMSD and Mg²⁺ correspondence were recalculated after this local fit. Comparison of the global-frame and locally fitted values was used to determine whether active-site agreement was already present in the overall complex alignment or resulted primarily from local optimization.

For visualization of global geometric agreement, the predicted *Nozomi* protein was coloured according to residue-wise Cα displacement from 6PR5. Predicted DNA was coloured according to the distance between each nucleotide backbone position and the nearest nucleotide-backbone position in the experimental complex; this DNA colouring was used as a geometric visualization rather than as a formal nucleotide alignment. The 6PR5 protein–DNA complex was displayed in semi-transparent grey.

### Identification and comparative analysis of *Nozomi* homologs

Putative *Nozomi* homologs were identified by searching the NCBI whole-genome shotgun sequence database with the predicted *Nozomi* transposase amino acid sequence using TBLASTN^64^. Protein sequences derived from selected homologous loci were compared with the *D. melanogaster Nozomi* transposase by pairwise global alignment using EMBOSS Needle^65^. Amino acid identity and similarity values were obtained from the resulting pairwise alignments. Transposase sequences were aligned using MUSCLE5^66^. The protein alignment was imported into R using the Biostrings package, and sequence names and ordering were standardized before downstream analysis. Amino acid identity between each transposase and the *D. melanogaster Nozomi* transposase was calculated from the multiple sequence alignment as the proportion of identical residues among positions at which both sequences contained an amino acid; alignment positions containing a gap in either sequence were excluded. The alignment and site-wise conservation were visualized using the ggmsa^67^ R package.

For focused comparison of the conserved catalytic region, amino acid identity was additionally calculated over alignment columns 292–437 using the same gap-exclusion procedure. This region was displayed together with an alignment conservation profile using ggmsa.

### TE decay index

To quantify degeneration of TE copies while accounting for both sequence divergence and sequence loss, we defined a TE decay index relative to a representative full-length reference element. TE sequences were aligned using `AlignSeqs` from DECIPHER, and pairwise nucleotide divergence from the reference was calculated under the Kimura two-parameter (K80) model using `dist.dna` from the ape R package, with pairwise deletion of alignment gaps.

For each TE copy, sequence coverage was calculated as: 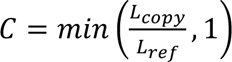, where *L_copy_* and *L_ref_* are the copy and reference lengths, respectively. K80 divergence was normalized as: 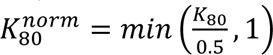. The TE decay index was then calculated as 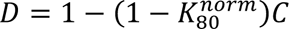. Thus, an intact sequence identical to the reference has *D* = 0, whereas increasing nucleotide divergence and/or sequence loss progressively increases the index toward 1.

## Data and code availability

All custom scripts used for the analyses in this study, together with FASTA files containing the nucleotide sequences of *Nozomi*, *Kodama* and related *Transib* elements and the amino acid sequences of their predicted transposases, are available at GitHub: https://github.com/foriin/Nozomi

## Acknowledgements

We thank Wolfgang Miller and Yuri Shevelyov for insightful discussions and valuable comments on the manuscript. The computations were enabled by resources provided by the National Academic Infrastructure for Supercomputing in Sweden (NAISS), partially funded by the Swedish Research Council through grant agreement no. 2022-06725. Some of the computing for this project was performed on the GenomeDK cluster. We would like to thank GenomeDK and Peter Ebert Andersen from Aarhus University for providing access to computational resources.

## Author contributions

A.I. performed data analysis and wrote the original draft with input from M.M. M.M. acquired funding.

---

**Supplementary Figure 1.** (A) (B) Copy-number comparison of *P-element* and *Transib1* insertions across DSPR and DrosEU long-read genome panels, respectively. Top: insertions classified as diverged partial, diverged full-length, non-diverged partial and non-diverged full-length copies. Bottom: non-diverged full-length insertions only. Insertions were classified as diverged when they differed from the consensus sequence by more than 10%.

(C) Genomic annotation of *Hopper*/*Kodama* insertions in dm6, A6 and BL2969 genomes relative to gene features.

(D) Genome browser shot showing an inversion on chr3R in the A6 genome relative to Canton-S genome flanked by a *Kodama* insertion. The inversion shown as the red rectangle on top.

**Supplementary Figure 2.** (A) Dot plot comparing the nucleotide sequences of *Nozomi* and *Kodama*.

(B) Heatmap showing pairwise nucleotide identity comparisons (%) between *Nozomi* insertions in A6 genome, including degenerated copies present on chromosome 3L designated as chr3L_fixed.

(C) Karyoplots showing the coordinates of *Nozomi* insertions in the A6 and BL2969 genomes after liftover to the dm6 genome assembly together with the coordinates of conserved piRNA clusters.

(D) Distribution of *Nozomi* and *Kodama* insertions in dm6 and the DSPR founder genomes relative to annotated gene features.

(E) Principal coordinate analysis of *Kodama* sequence diversity across three D. melanogaster genomes. PCoA was based on alignment-normalized pairwise Hamming distances among *Kodama* sequences from dm6, A6, and BL2969. Points are coloured by genome. Point size represents the number of identical copies corresponding to each collapsed A6 or BL2969 sequence; dm6 sequences are shown at a fixed size. Axis labels indicate the percentage of variation explained by each principal coordinate.

(F) Relationship between pairwise genomic distance and K80 nucleotide divergence among *Kodama* insertions. Each point represents a pair of insertions, and the fitted trend line summarizes the overall association between genomic proximity and sequence divergence.

**Supplementary Figure 3.** (A) Coverage of Pool-seq reads from 48 geographically diverse *D. melanogaster* populations mapped to the autonomous *Nozomi* consensus. Coverage was first normalized on the number of reads mapped to dm6 genome (reads per million dm6 mapped reads, RPM) and then on the maximum RPM value.

B) Histograms showing length of *Transib-6* insertions in *D. bipectinata* genome. Top: all identified insertions. Bottom: insertions with at least first 8 nt of left and right TIRs preserved. Dashed line demarcates the median insertion length.

**Supplementary Figure 4.** Multiple sequence alignments of collapsed *Kodama* and *Nozomi* sequences from the A6 (A) and BL2969 (B) genomes, focused on the boundaries of the characteristic 1,380-bp central deletion. Thirty-five alignment positions flanking each deletion boundary and 15 positions from each edge of the deleted region are shown; the intervening portion of the deletion is omitted and indicated by a double slash. Each row represents a distinct sequence, with its length and the number of identical copies represented by that sequence indicated on the left. Nucleotides are colored by base identity, and gaps are left blank.

**Supplementary Figure 5.** (A) Multiple-sequence alignment of the catalytic core of *Transib* transposases analyzed in this study. Three alignment blocks centred on the conserved DDE catalytic triad are shown. Red asterisks mark the catalytic residues, and numbers in parentheses indicate the number of amino acids omitted from each sequence between adjacent blocks. Amino acids are coloured according to the Clustal scheme.

(B) Amino acid identity of *Nozomi* homologs’ RNAse-H fold in the catalytic domain.

(C) Coverage of whole-genome sequencing reads from geographically diverse *D. suzukii* populations mapped to the autonomous *Hayabusa* consensus, showing continuous coverage across the element and no evidence of the fragmented pattern observed for *Nozomi* in *D. melanogaster*.

(D) Distribution of K80 distances of *Kodama* insertions to *Nozomi* sequence in dm6, A6 and BL2969 genomes.

(E) Distribution of K80 distances to the corresponding consensus sequences for *Hayabusa* and *Darjeeling* copies in the *D. suzukii* genome and *Nozomi* copies in the *D. melanogaster* dm6 genome.

