## Supplementary Figures for "The Little Transib That Could – and Could Not: Contrasting Invasion Outcomes within a Novel DDE DNA Transposon Lineage"

### Supplementary figure 1

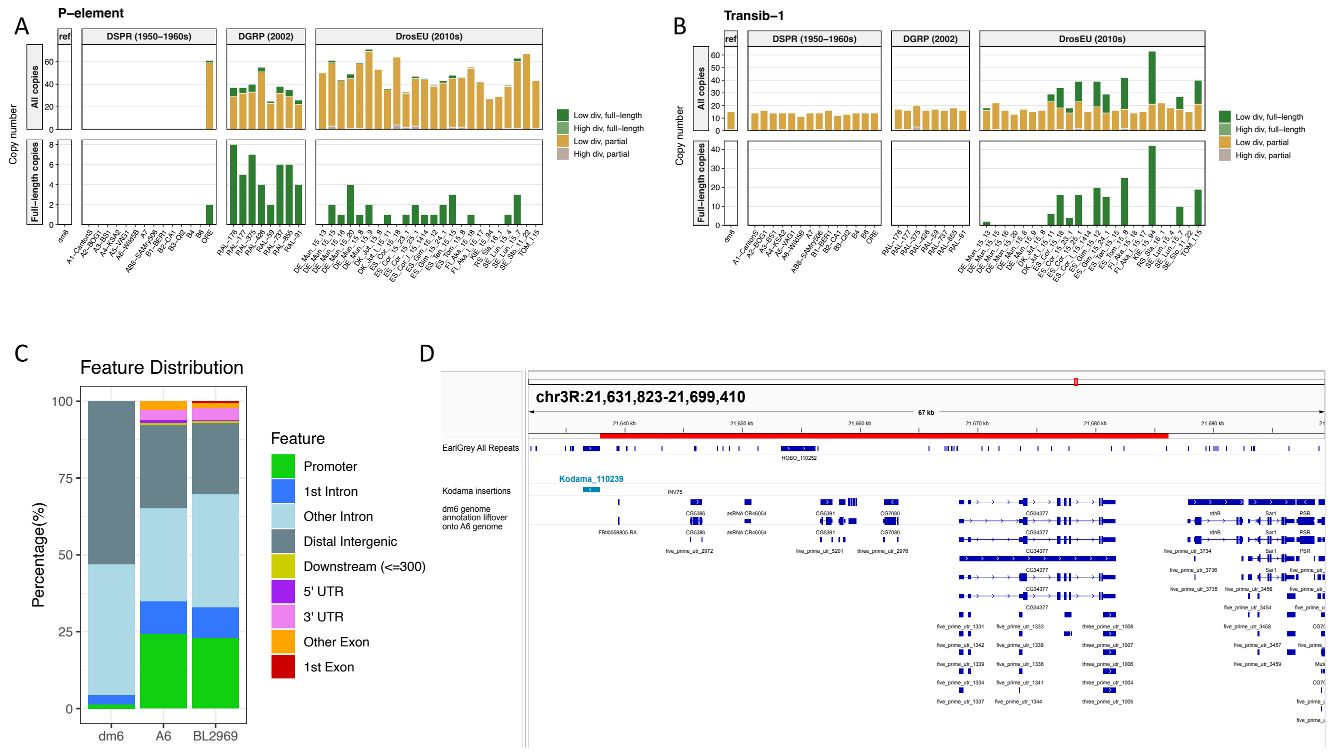

### Supplementary figure 2

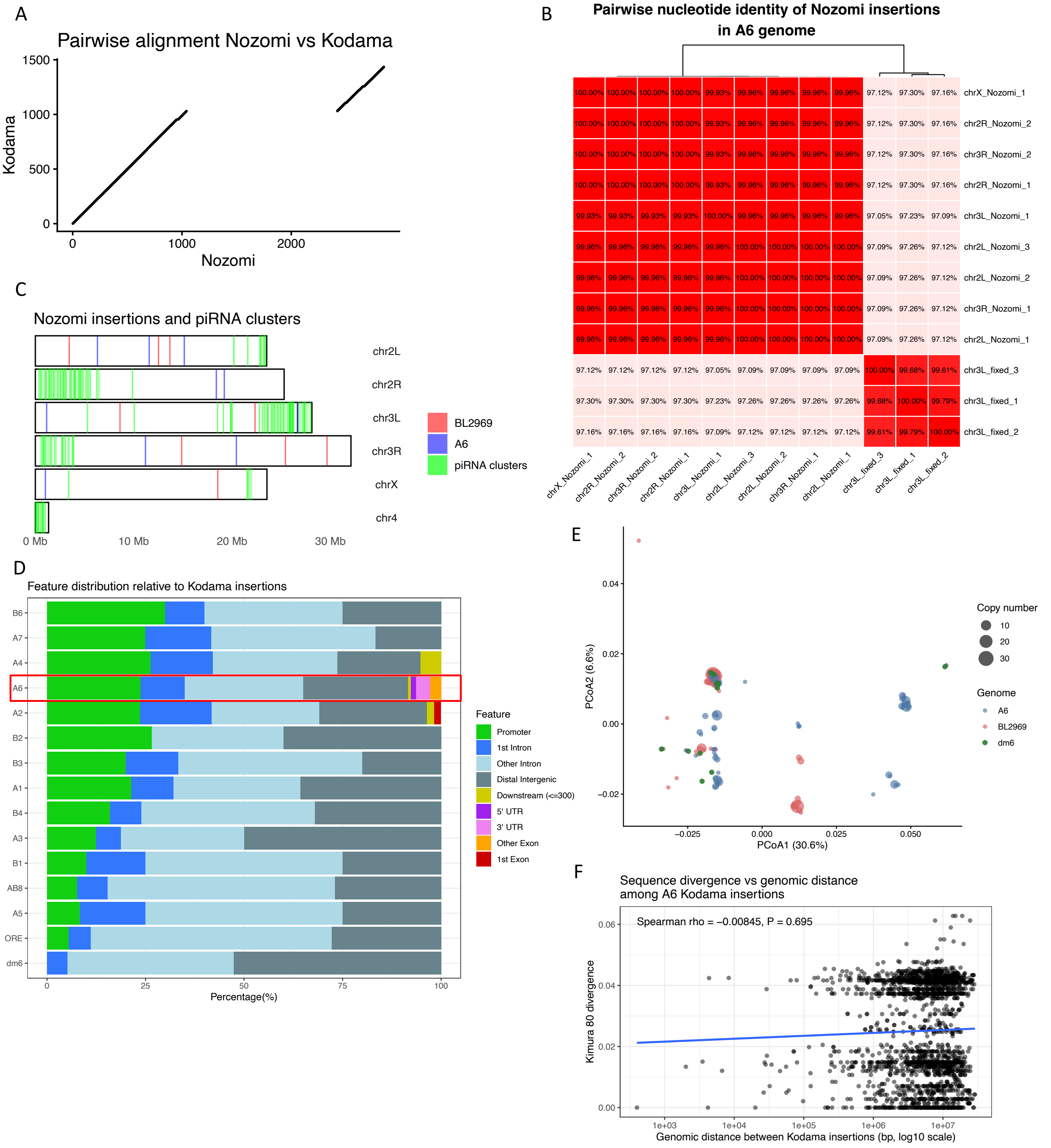

### Supplementary figure 3

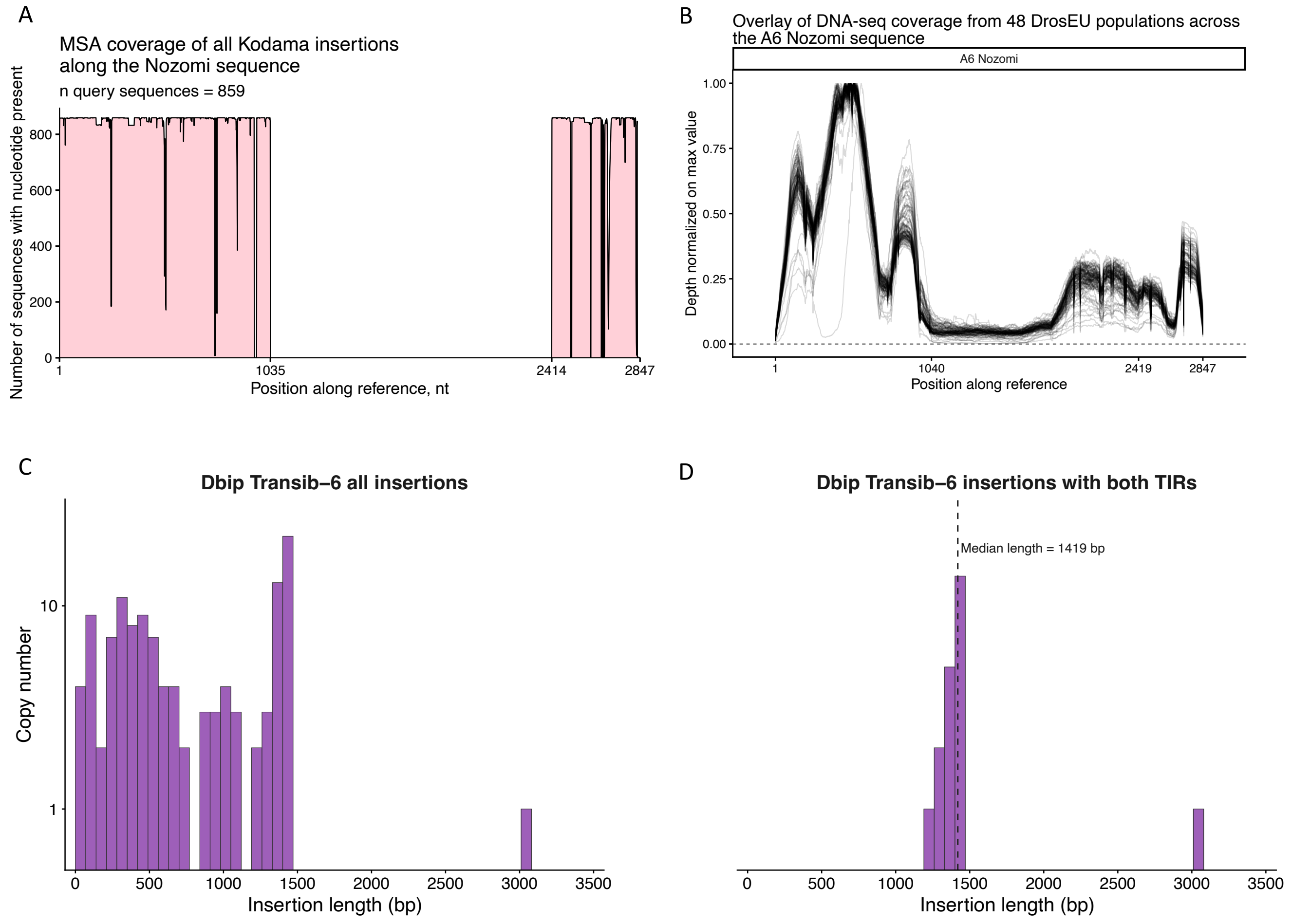

### Supplementary figure 4

A

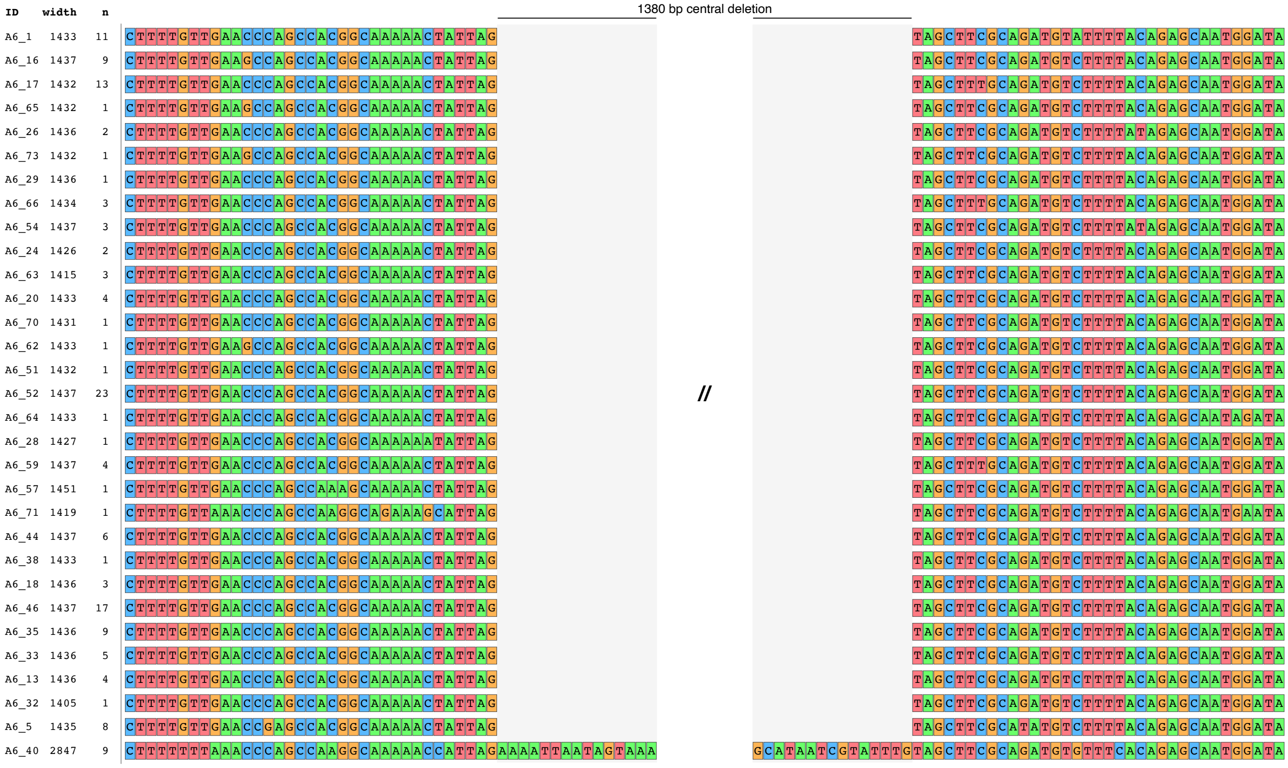


B

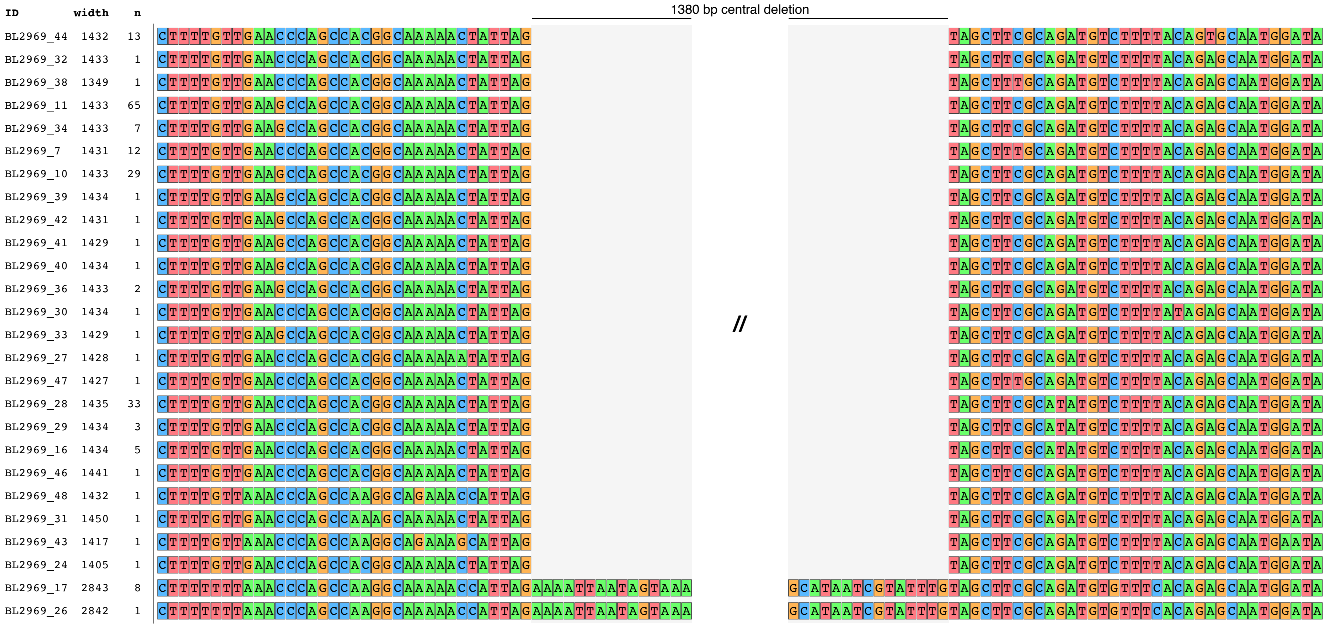

### Supplementary figure 5

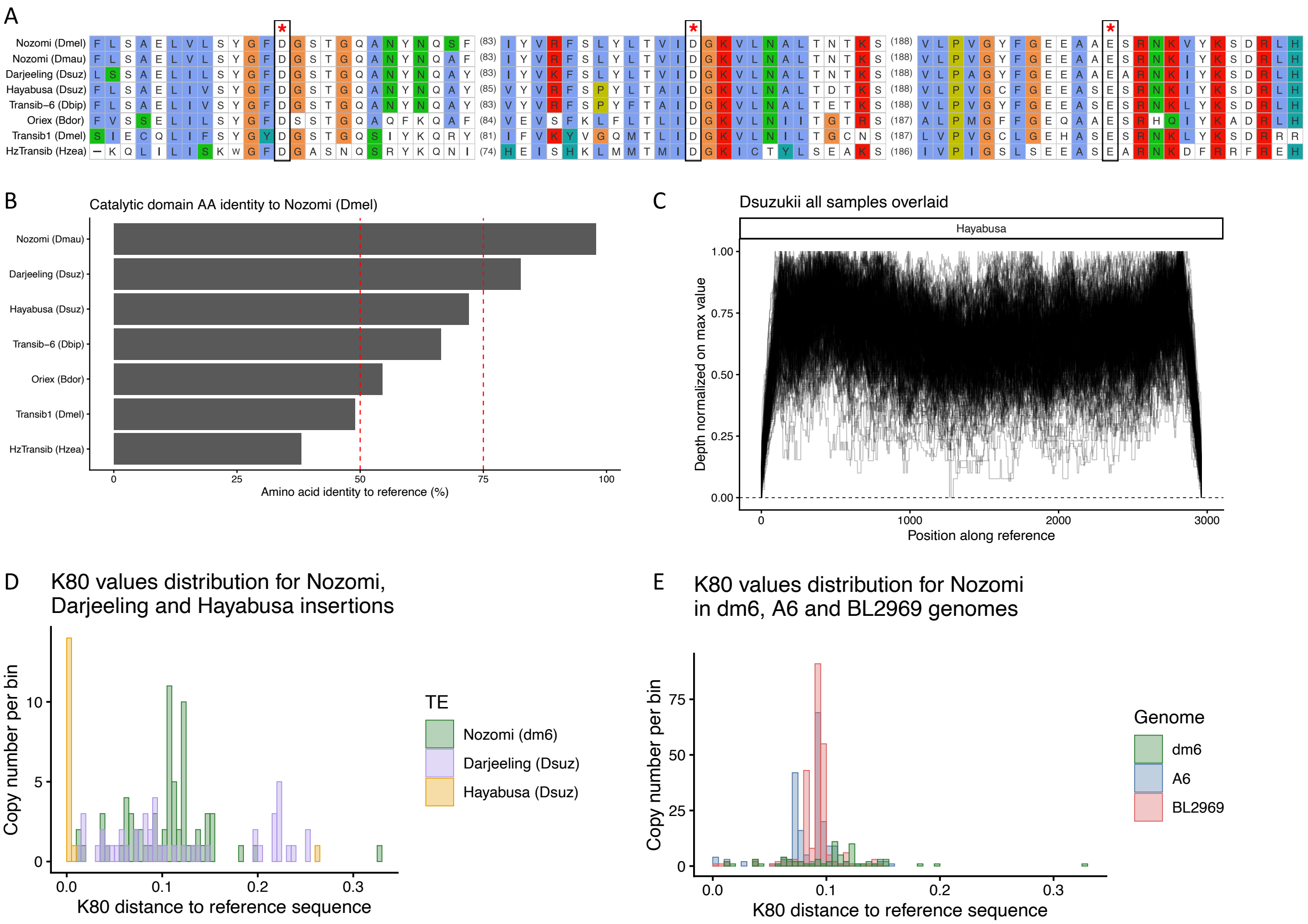
